# Estimating muscle activation from EMG using deep learning-based dynamical systems models

**DOI:** 10.1101/2021.12.01.470827

**Authors:** Lahiru N. Wimalasena, Jonas F. Braun, Mohammad Reza Keshtkaran, David Hofmann, Juan Álvaro Gallego, Cristiano Alessandro, Matthew C. Tresch, Lee E. Miller, Chethan Pandarinath

**Affiliations:** Wallace H. Coulter Department of Biomedical Engineering, Emory University and Georgia Institute of Technology, Atlanta, GA, USA; Department of Electrical and Computer Engineering, Technical University of Munich, Munich, Germany; School of Life Sciences, École Polytechnique Fédérale de Lausanne, Lausanne, Switzerland; Department of Physics, Emory University, Atlanta, GA, USA; Initiative in Theory and Modeling of Living Systems, Emory University, Atlanta, GA, USA; Department of Bioengineering, Imperial College London, London, UK; Department of Physiology, Northwestern University, Chicago, IL, USA; Department of Brain and Behavioral Sciences, University of Pavia, Pavia, Italy; School of Medicine and Surgery, University of Milano-Bicocca; Department of Biomedical Engineering, Northwestern University, Evanston, IL, USA; Shirley Ryan AbilityLab, Chicago, IL, USA; Department of Physical Medicine and Rehabilitation, Northwestern University, Chicago, IL, USA; Department of Neurosurgery, Emory University, Atlanta, GA, USA

**Keywords:** EMG, deep learning, dynamical systems, motor control

## Abstract

**Objective:** To study the neural control of movement, it is often necessary to estimate how muscles are activated across a variety of behavioral conditions. However, estimating the latent command signal that underlies muscle activation is challenging due to its complex relation with recorded electromyographic (EMG) signals. Common approaches estimate muscle activation independently for each channel or require manual tuning of model hyperparameters to optimally preserve behaviorally-relevant features.

**Approach:** Here, we adapted AutoLFADS, a large-scale, unsupervised deep learning approach originally designed to de-noise cortical spiking data, to estimate muscle activation from multi-muscle EMG signals. AutoLFADS uses recurrent neural networks (RNNs) to model the spatial and temporal regularities that underlie multi-muscle activation.

**Main Results:** We first tested AutoLFADS on muscle activity from the rat hindlimb during locomotion, and found that it dynamically adjusts its frequency response characteristics across different phases of behavior. The model produced single-trial estimates of muscle activation that improved prediction of joint kinematics as compared to low-pass or Bayesian filtering. We also tested the generality of the approach by applying AutoLFADS to monkey forearm muscle activity from an isometric task. AutoLFADS uncovered previously uncharacterized high-frequency oscillations in the EMG that enhanced the correlation with measured force compared to low-pass or Bayesian filtering. The AutoLFADS-inferred estimates of muscle activation were also more closely correlated with simultaneously-recorded motor cortical activity than other tested approaches.

**Significance:** Ultimately, this method leverages both dynamical systems modeling and artificial neural networks to provide estimates of muscle activation for multiple muscles that can be used for further studies of multi-muscle coordination and its control by upstream brain areas.

## Introduction

Executing movements often requires activating muscles distributed across the body in precisely coordinated time-varying patterns. Understanding how the nervous system selects and generates these patterns remains a central goal of motor neuroscience. Critical to this effort is obtaining estimates of the neural commands to muscles and evaluating how they are related both to task demands and to neural activity in motor related brain areas. However, a key barrier in this evaluation is the difficulty of estimating muscle activation from electromyographic (EMG) recordings.

EMG recordings are typically rectified and then further processed to obtain an estimate of the activation envelope for a given muscle. However, processing this rectified signal to extract muscle activation is non-trivial. A wide variety of filtering approaches have been proposed, ranging from a simple linear, low-pass filter (Clancy et al., 2001; D’Alessio and Conforto, 2001; Hogan and Mann, 1980) to nonlinear, adaptive Bayesian filtering approaches that aim to model the time-varying statistics of the EMG (Hofmann et al., 2016; Sanger, 2007). Although these are reasonable approaches, determining the optimal parameters for these filters is challenging, and typically results in arbitrary or non-optimal choices. Some parameter combinations may result in less smoothing that leads to more variable signals that are difficult to interpret, while other combinations may suppress high frequency features that may be relevant to behavior. Another common approach is to simplify signals further by averaging across repeats of the same behavior, which destroys any correspondence with across-trial behavioral variation.

In cases where EMG is simultaneously recorded from multiple muscles, many filtering approaches treat multi-muscle EMG recordings as independent signals. These approaches fail to leverage potential information in the coordinated activation of muscles. Previous work in the field of muscle synergies has demonstrated underlying spatial and temporal regularities in multi-muscle activation across many model systems, suggesting that co-variation patterns across muscles may provide additional information for the estimation of muscle activations (Tresch and Jarc, 2009).

Here we adapted a large-scale deep learning approach, AutoLFADS, leveraging advances in deep generative modeling to estimate muscle activation from multi-muscle recordings. In particular, AutoLFADS models the dynamics (i.e., spatial and temporal regularities) underlying multi-muscle activity using recurrent neural networks (RNNs). The large-scale optimization approach allows AutoLFADS the flexibility to be trained out-of-the-box, i.e., without manual adjustment for different datasets.

A separate class of methods to those discussed here aims to use EMG and other signals, as well as musculoskeletal parameters, to estimate activation torques, joint torques and joint kinematics, with potential applications to musculoskeletal simulation and prosthetic device control (Nasr et al., 2021). However, such methods have a different focus – aside from their system inputs, there is no subsequent representation of muscle activation. In contrast, our approach uses simultaneously-recorded EMG from multiple muscles to make improved estimates of muscle activation itself. While our approach could be used as a preprocessing strategy to support classification and control, its primary goals are to produce a high-quality estimate of the underlying muscle activation signal(s), appropriate for understanding basic motor control properties of the brain. As such, we extend a large body of previous literature - particularly, linear filtering or Bayesian filtering - which have largely approached this problem by looking at the activity from a single muscle at a time (Clancy et al., 2001; Hofmann et al., 2016; Sanger, 2007). The key advantages of our approach are accounting for the known spatial and temporal regularities underlying EMG signals - as highlighted by the field of muscle synergies, for example (Tresch and Jarc, 2009) - and the use of powerful deep learning methods to uncover these coordinated patterns on fast timescales and on individual trials. In previous work, LFADS was used to model the dynamics of the brain area under study. In this case, muscle activity is hypothesized to be the output of a dynamical system that we cannot directly observe via EMG - i.e., spinal circuitry and inputs from the many different brain areas that play a role in activating muscles. Given this lack of access, the key question was whether we would see improvements in the quality of the estimates of muscle activation by modeling EMG as the output of a dynamical system.

We first applied AutoLFADS to multi-muscle recordings in rat hindlimb during locomotion. We found that AutoLFADS functions adaptively to adjust its frequency response characteristics dynamically according to the time course of different phases of behavior. As with other latent variable models (Hofmann et al., 2016; Sanger, 2007), we validated our approach by testing and comparing how well the representations inferred by AutoLFADS correlated with behavioral output. We showed that AutoLFADS enables more accurate single-trial joint angular acceleration predictions from EMG than the predictions of standard filtering techniques (i.e., smoothing, Bayesian filtering). We next applied AutoLFADS to multi-muscle recordings in monkey forearm during an isometric force-control task. Unexpectedly, AutoLFADS identified previously uncharacterized 10-50 Hz oscillations shared across flexor muscles, and resulted in enhanced coherence with recorded force signals at higher frequencies. Finally, the AutoLFADS-inferred muscle activations corresponded closely to simultaneously recorded activity in the motor cortex, suggesting that the model preserves correlations with both descending motor commands and behavior. These results show that AutoLFADS reveals subtle features in muscle activity that may otherwise be overlooked or eliminated, and thus may allow new insights into motor control by elucidating relationships between muscle activation signals in the brain and spinal cord with high temporal precision on a single-trial level.

## Results

### Adapting AutoLFADS to model dynamics underlying EMG activity

We begin with a model schematic outlining the adapted AutoLFADS approach (**Fig. 1**). **m**[t] is the time-varying vector of rectified multi-muscle EMG. Our goal is to find 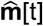, an estimate of muscle activation from **m**[t] that preserves the neural command signal embedded in the recorded EMG activity. Common approaches to estimating 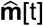 typically involve applying a filter to each EMG channel independently, ranging from a simple linear smoothing filter to a nonlinear, adaptive Bayesian filter. While these approaches effectively reduce high-frequency noise, they have two major limitations. First, they require supervision to choose hyperparameters that affect the output of the filters. For example, using a smoothing (low-pass) filter requires a choice of a cutoff frequency and order, which sets an arbitrary limit of the EMG frequency spectrum that is considered “signal”. Setting a lower cutoff yields smoother representations with less high-frequency content; however, this may also come at the expense of eliminating high-frequency activation patterns that may be behaviorally-relevant. Second, by filtering muscles independently, standard smoothing approaches fail to take advantage of potential information in the shared structure across muscles. In particular, provided that muscles are not activated independently (Tresch and Jarc, 2009), the estimate of a given muscle’s activation can be improved by having access to recordings from other muscles.

**Fig. 1.**
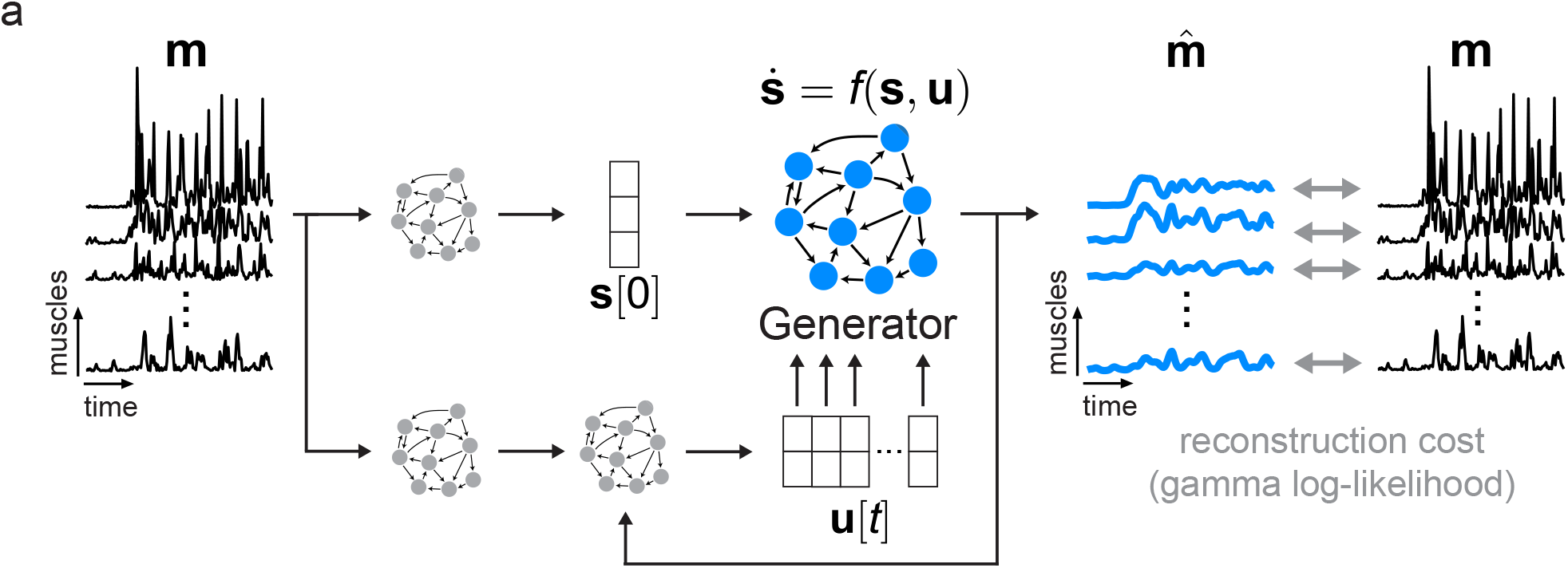
Modeling dynamics underlying EMG activity by adapting AutoLFADS. Model schematic. We modified AutoLFADS to accommodate the statistics of EMG activity using a gamma observation model rather than Poisson. **m** is a small window from a continuous recording of rectified EMG from a group of muscles. **m**[t] is a vector representing the multi-muscle rectified EMG at each timepoint. 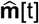 is an estimate of muscle activation from the EMG activation envelopes **m**[t]. Our approach models 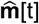 as the output of a nonlinear dynamical system with shared inputs to the muscles. The Generator RNN is trained to model the state update function *f*, which captures the spatial and temporal regularities that underlie the coordinated activation of multiple muscles. The other RNNs provide the Generator with an estimate of initial state **s**[0] and a time-varying input **u**[t], allowing the dynamical system to evolve in time to generate **s**[t]. 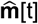 is taken as a nonlinear transformation of **s**[t]. All of the model’s parameters are updated via backpropagation of the gamma log-likelihood of observing the input rectified EMG data.

Our approach overcomes the limitations of standard filtering by leveraging the shared structure and potential nonlinear dynamics describing the time-varying patterns underlying **m**[t]. We implemented this approach by adapting latent factor analysis via dynamical systems (LFADS), a deep learning tool previously developed to study neuronal population dynamics (**Fig. 1**) (Pandarinath et al., 2018). Briefly, LFADS comprises a series of RNNs that model observed data as the noisy output of an underlying nonlinear dynamical system. The internal state of one of these RNNs, termed the *Generator*, explicitly models the evolution of **s**[t] i.e., the state of the underlying dynamical system. The Generator is trained to model *f* i.e., the state update function, by tuning the parameters of its recurrent connections to model the spatial and temporal regularities governing the coordinated time-varying changes of the observed data. The Generator does not have access to the observed data so it has no knowledge of the identity of recorded muscles or the observed behavior. Instead, the Generator is only provided with an initial state **s**[0], and a low-dimensional, time-varying input **u**[t] to model the progression of **s**[t] as a discrete-time dynamical system. The model then generates 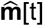, where **m**[t] is taken as a sample of 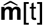 through a gamma observation model (Details are provided in *Methods*). The model has a variety of hyperparameters that control training and prevent overfitting. These hyperparameters are automatically tuned using the AutoLFADS large-scale optimization framework we previously developed (Keshtkaran et al., 2021; Keshtkaran and Pandarinath, 2019).

While the core LFADS model remains the same in its application for EMG activity as for cortical activity, we did not have prior guarantee that adapting the observation model alone would be sufficient to provide us with good estimates of muscle activation from EMG. As a key distinction to previous applications of LFADS, we used data augmentation during training. In our initial modeling, we found that models could overfit to high-magnitude events that occurred simultaneously across multiple channels. To mitigate this issue, we implemented a data augmentation strategy (“temporal shift”) during model training to prevent the network from drawing a direct correspondence between specific values of the rectified EMG used both as the input to the model and at the output to compute the reconstruction cost (details provided in *Methods*).

### Applying AutoLFADS to EMG from locomotion

We first tested the adapted AutoLFADS approach using data recorded from the right hindlimb of a rat walking at constant speed on a treadmill (Alessandro et al., 2018). Seven markers were tracked via high-speed video, which allowed the 3 joint angular kinematic parameters (hip, knee, and ankle) to be inferred, along with the time of foot contact with the ground (i.e., foot strike) and the time when the foot left the ground (i.e., foot off) for each stride (**Fig. 2a**). In parallel, EMG signals were recorded from 12 hindlimb muscles using intramuscular electrodes. Continuous hindlimb EMG recordings were segmented using behavioral event annotations, considering foot strike as the beginning of each stride. Aligning estimates to behavioral events allowed us to decompose individual strides into the “stance” phase, where the foot is in contact with the ground, and the “swing” phase, where the foot is in the air in preparation for the next stride.

**Fig. 2.**
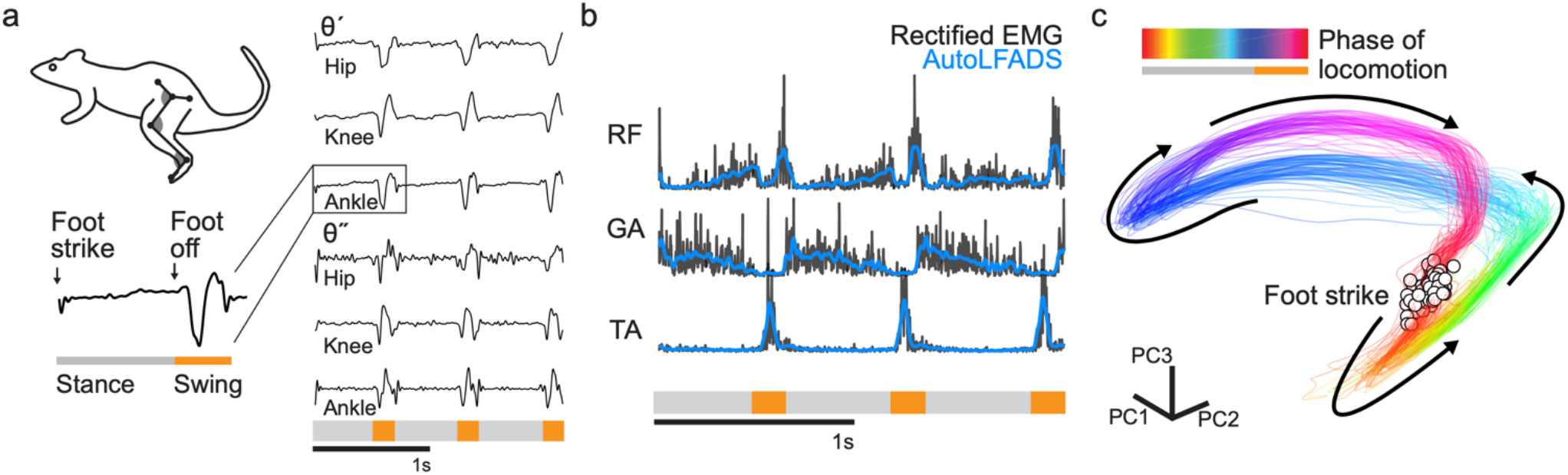
Applying AutoLFADS to rat locomotion. (a) Schematic of rat locomotion. Examples of three joint angular velocities (θ’) and angular accelerations (θ”) are shown for three consecutive strides (derived from recorded joint marker position tracking). Zoomed segment (left) indicates periods of “stance” and “swing” during a stride. The swing phase of each stride is labeled below in orange. (b) Example AutoLFADS output (blue) overlaid on rectified, unfiltered EMG (black) for three muscles in the rat hindlimb (RF: rectus femoris, GA: gastrocnemius, TA: tibialis anterior). (c) Top 3 principal components of the AutoLFADS output for individual strides, colored by phase of locomotion (cumulative variance explained: 92%). Arrows indicate the direction of the state evolution over time. Foot strike events are labeled with white dots.

We trained AutoLFADS models on EMG activity from the rat hindlimb during locomotion to estimate the muscle activation for each muscle (**Fig. 2b**). AutoLFADS output substantially reduced high-frequency fluctuations while still tracking fast changes observed in the rectified EMG, reflecting both smooth phases of the behavior (e.g., the stance phase) and rapid behavioral changes (e.g., the beginning of the swing phase). AutoLFADS models were trained in a completely unsupervised manner, which meant the models did not have information about the behavior, the identity of the recorded muscles, or any additional information outside of the rectified multi-muscle EMG activity. Visualizing muscle activity using state space representations helps to illustrate how AutoLFADS was capable of modeling dynamics. We performed dimensionality reduction via principal components analysis to reduce the 12 AutoLFADS output channels to the top three components (**Fig. 2c**). AutoLFADS estimates from individual strides followed a stereotyped trajectory through state space: state estimates were tightly clustered at the time of foot strike, and evolved in a manner where activity at each phase of the gait cycle occupied a distinct region of state space.

### AutoLFADS adapts filtering to preserve behaviorally relevant, high-frequency features

A key feature that differentiates AutoLFADS from previous approaches is the ability to dynamically adapt its filtering properties to phases of behavior where signal content exists at different timescales. In contrast, approaches like low-pass filtering cannot adaptively determine what components are features of the muscle activation; instead, they apply a fixed cutoff frequency throughout the signal, regardless of the potential changes in timescale that may occur during different phases of behavior. We demonstrated the adaptive nature of the AutoLFADS model by comparing its filtering properties during the swing and stance phases of locomotion. Swing phase contains higher-frequency transient features related to foot lift off and touch down, which are in contrast to the lower frequency changes in muscle activity during the stance phase. We computed the power spectral density (PSD) for the AutoLFADS inferred muscle activations, rectified EMG smoothed with a 20 Hz low pass filter, and the unfiltered rectified EMG within 350 ms windows centered on the stance and swing phases of locomotion. We then used these PSD estimates to compute the frequency responses of the AutoLFADS model - i.e., approximating it as a linear filter - and of the 20 Hz low pass filter over many repeated strides within the two phases (see Methods).

As expected, the 20 Hz low pass filter had a consistent frequency response during the swing and stance phases (**Fig. 3**). In contrast, the AutoLFADS output resulted in substantially different frequency response estimates that enhanced high frequencies during the rapid changes of the swing phase and attenuated them during stance. Less obviously, the frequency responses estimated for AutoLFADS differed across muscles, despite its reliance on a shared dynamical system. Some AutoLFADS frequency responses (e.g., GS and RF) changed substantially between stance and swing in the 10-20 Hz frequency band; however, other muscles (e.g., IL and GA) shifted only slightly.

**Fig. 3.**
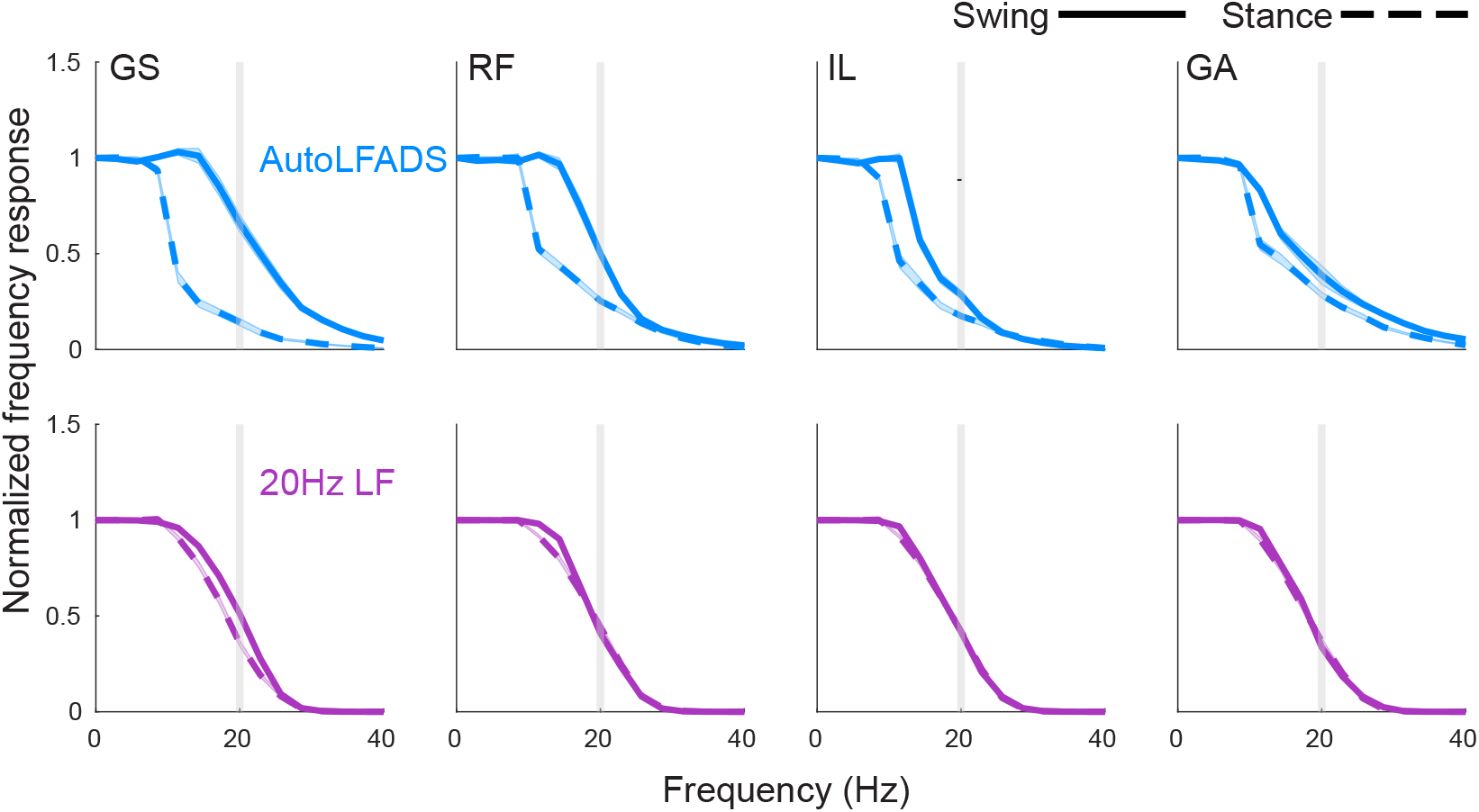
AutoLFADS filtering adapts to preserve behaviorally relevant, high-frequency EMG features, in contrast to static low-pass filters. (a) Approximated frequency responses of lf_20_ EMG (purple) and AutoLFADS (blue) for 4 muscles in the swing (solid) and stance (dashed) phases (GS: gluteus superficialis, RF: rectus femoris, IL: iliopsoas, GA: gastrocnemius). Shaded area represents SEM of estimated frequency responses from single stride estimates. Estimated frequency responses from single strides were fairly consistent resulting in small SEM. Gray vertical line at 20Hz signifies low pass filter cutoff.

### AutoLFADS is more predictive of behavior than other filtering approaches

Training AutoLFADS models in an unsupervised manner (i.e., without any information about simultaneously recorded behavior, experiment cues, etc.) raises the question of whether the muscle activation estimates found by modeling multi-muscle dynamics were informative about behavior. To assess this, we compared muscle activations estimated by AutoLFADS to those estimated using standard filtering approaches, by evaluating how well each estimate could decode simultaneously recorded behavioral variables (**Fig. 4**). We performed single-timestep optimal linear estimation (OLE) on joint angular accelerations (**Fig. 4a**) that captured high-frequency changes in the joint angular kinematics of rats during locomotion.

**Fig. 4.**
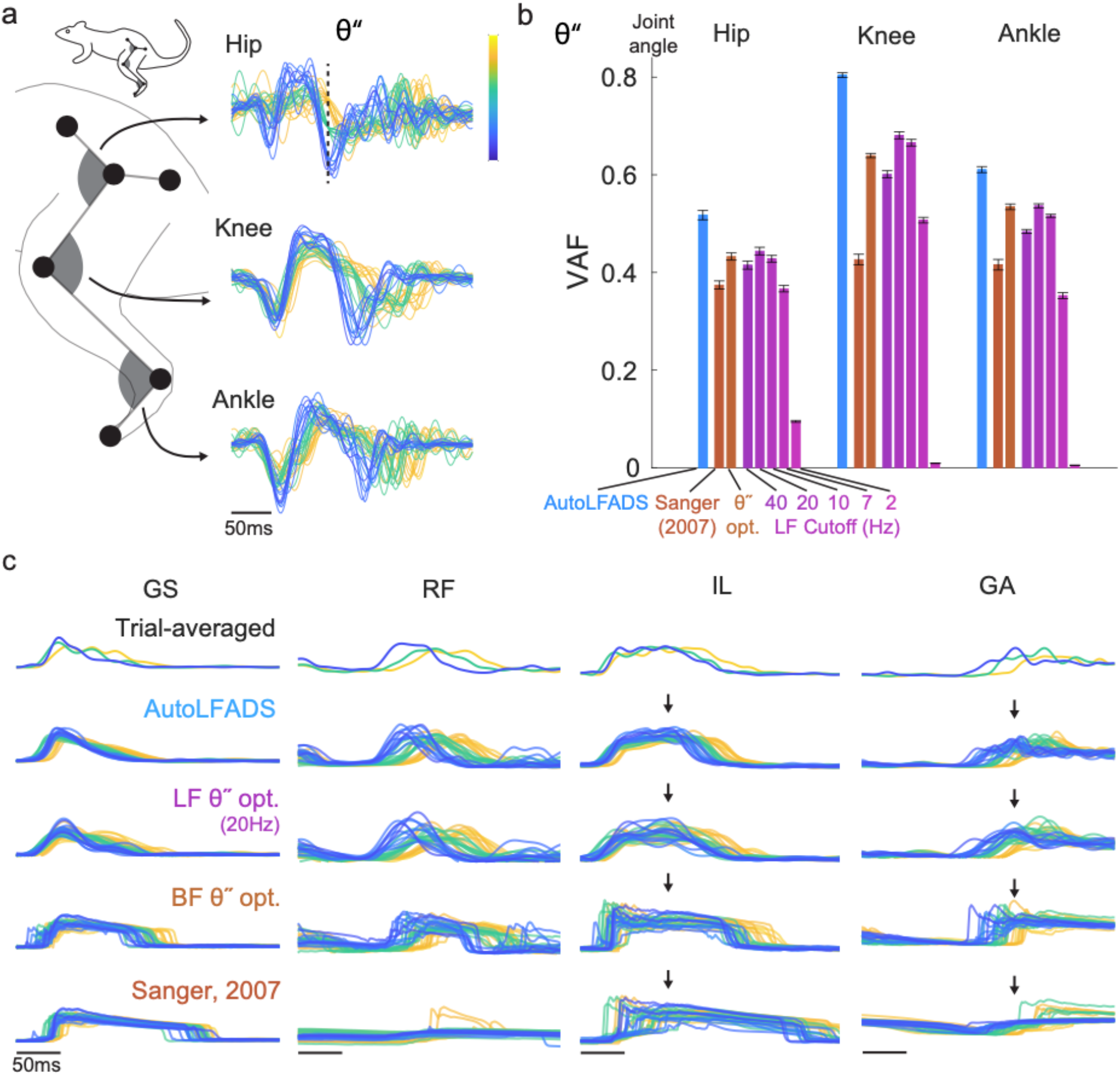
AutoLFADS muscle activation estimates are more predictive of joint angular acceleration than other filtering approaches. (a) Visualization of single trial joint angular accelerations (θ”) for hip, knee, and ankle joint angular kinematic parameters. Trials were aligned to the time of foot off using an optimal shift alignment method. Trials were grouped and colored based on the magnitude of the hip angular acceleration at a specific time (indicated by dashed line). (b) Joint angular acceleration decoding performance for AutoLFADS, Bayesian filtering approaches, and multiple cutoff frequencies of low-pass filters, quantified using variance accounted for (VAF). Each bar represents the mean +/− SEM across cross-validated prediction VAF values for a given joint. (c) Visualization of single trial muscle activation estimates for AutoLFADS, EMG smoothed with a 20 Hz low-pass filter (LF θ” opt.), Bayesian filtering with hyperparameters optimized to maximize prediction of θ” (BF θ” opt.), and Bayesian filtering using the same hyperparameters as Sanger, 2007. Alignment and coloring of single trials are the same as (a). Arrows highlight visual features that differentiate muscle activation estimates from tested methods.

AutoLFADS significantly outperformed all low-pass filter cutoffs (ranging from 2 to 40Hz) and Bayesian filtering approaches for prediction of angular acceleration at the hip, knee, and ankle joints (differences in variance accounted for (VAF); hip: p=4.44e-07, knee: p=9.81e-09, ankle: p=2.86e-06 paired t-test; **Fig. 4b**). Visualizing the muscle activation estimates across individual strides underscored the differences between the approaches (**Fig. 4c**). Even Bayesian filters optimized to maximize prediction accuracy of joint angular accelerations did significantly worse than AutoLFADS, primarily because they estimated the activation offsets poorly. Even though the AutoLFADS model was trained in an unsupervised manner with no knowledge of the joint angular accelerations, our approach outperformed other approaches that had manually tuned hyperparameters to yield the best possible decoding performance.

### Applying AutoLFADS to EMG from isometric wrist tasks

To better understand the generalizability of our approach to other model systems, muscle groups, and behaviors, we also analyzed data from a monkey performing a 2-D isometric wrist task (**Fig. 5a**) (Gallego et al., 2020). EMG activity was recorded from 14 channels (including 12 forearm muscles) during a task requiring the monkey to generate torques isometrically about the wrist. The monkey’s forearm was restrained and its hand clamped securely in a box instrumented with a 6 DOF load cell. Brain activity was simultaneously recorded via a 96-channel microelectrode array in the hand area of contralateral M1. The monkeys controlled the position of a cursor displayed on a monitor by modulating the force they exerted. Flexion and extension forces moved the cursor right and left, respectively, while forces along the radial and ulnar deviation axis moved the cursor up and down. Monkeys controlled the on-screen cursor to acquire targets presented in one of eight locations in a center-out behavioral task paradigm. Each trial consisted of a step-like transition from a state of minimal muscle activity prior to target onset to one of eight phasic-tonic muscle activation patterns (i.e., one for each target) that enabled the monkey to hold the cursor at the target. These step-like changes and distinct muscle activation patterns across target locations contrasted the cyclic behavior and highly repetitive activation patterns observed during rat locomotion, allowing us to evaluate AutoLFADS in two fundamentally different contexts.

**Fig. 5.**
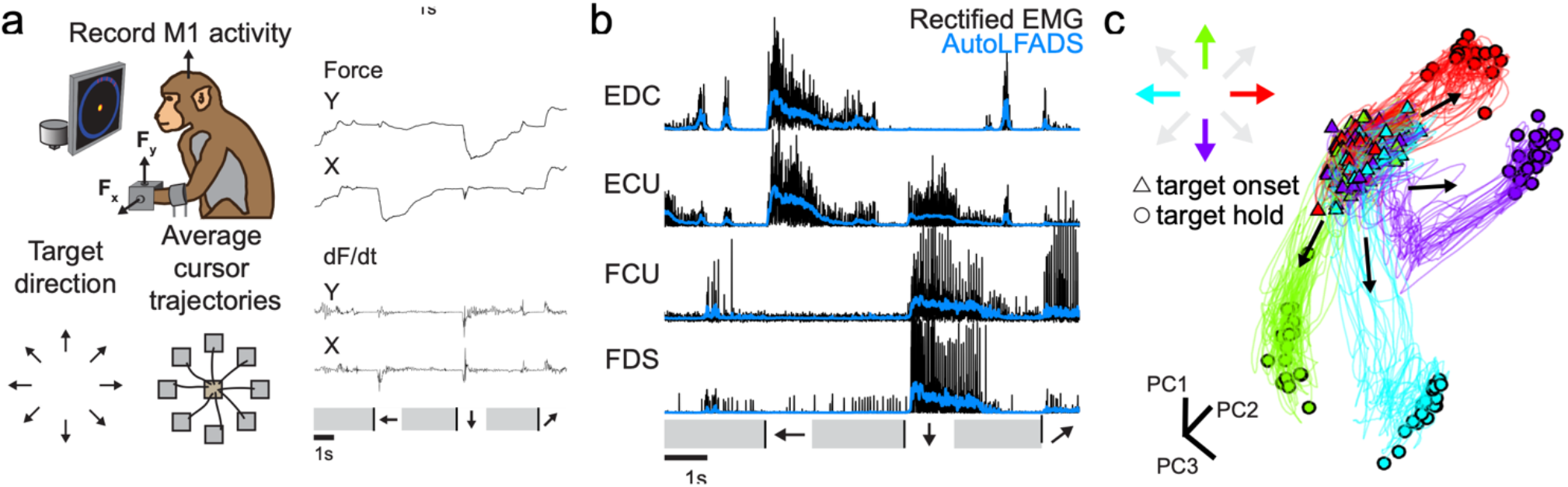
Applying AutoLFADS to isometric wrist tasks. (a) Schematic of monkey isometric wrist task. Force and dF/dt are shown for three consecutive successful trials. The target acquisition phases are indicated by arrows corresponding to the target directions. Black lines denote target onset. Grey intervals denote return to and hold within the central target. (b) Example AutoLFADS output (blue) overlaid on rectified EMG (black) for four forearm muscles with a moment about the wrist (EDC: extensor digitorum communis, ECU: extensor carpi ulnaris, FCU: flexor carpi ulnaris, FDS: flexor digitorum superficialis). (c) Top three principal components of AutoLFADS output for four conditions, colored by target hold location (cumulative variance explained: 88%).

We trained an AutoLFADS model on EMG activity from the monkey forearm during the wrist isometric task to estimate the muscle activation for each muscle (**Fig. 5b**). As for the locomotion data, the AutoLFADS output substantially reduced high-frequency fluctuations while still tracking changes in the rectified EMG, reflecting both smooth phases of the behavior (e.g., the period prior to target onset in the wrist task) and rapid behavioral changes (e.g., the target acquisition period in the wrist task). We again performed dimensionality reduction to reduce the AutoLFADS output (14 channels) to the top three PCs. In the monkey data, the muscle activation estimates from AutoLFADS began consistently in a similar region of state space at the start of each trial, then separated to different regions at the target hold period. This demonstrates that the model was capable of inferring distinct patterns of muscle activity for different target directions, even without the cyclic, repeated task structure of the locomotion data (**Fig. 5c**).

### AutoLFADS uncovers high-frequency oscillatory structure across muscles during isometric force production

An interesting attribute of the isometric wrist task is that for certain target conditions that require significant flexion of the wrist, the x component of force contains high-frequency oscillations – made evident by visualizing the dF/dt (**Fig. 6a**). Corresponding oscillatory structure can be seen in the rectified EMG of flexor muscles (**Fig. 6b**). We found that muscle activation estimates from the tested approaches preserve these high-frequency features from the rectified EMG differently (**Fig. 6c**) AutoLFADS output showed clear, consistent oscillations that lasted for 3-4 cycles, while other approaches such as Bayesian filtering (using the hyperparameters from Sanger, 2007) smoothed over these features. Oscillations in dF_x_/dt were visible for three of the eight target conditions (**Fig. 6d**) and corresponding oscillations were observed across the flexor muscles in the AutoLFADS output (**Fig. 6e**) in those trials.

**Fig. 6.**
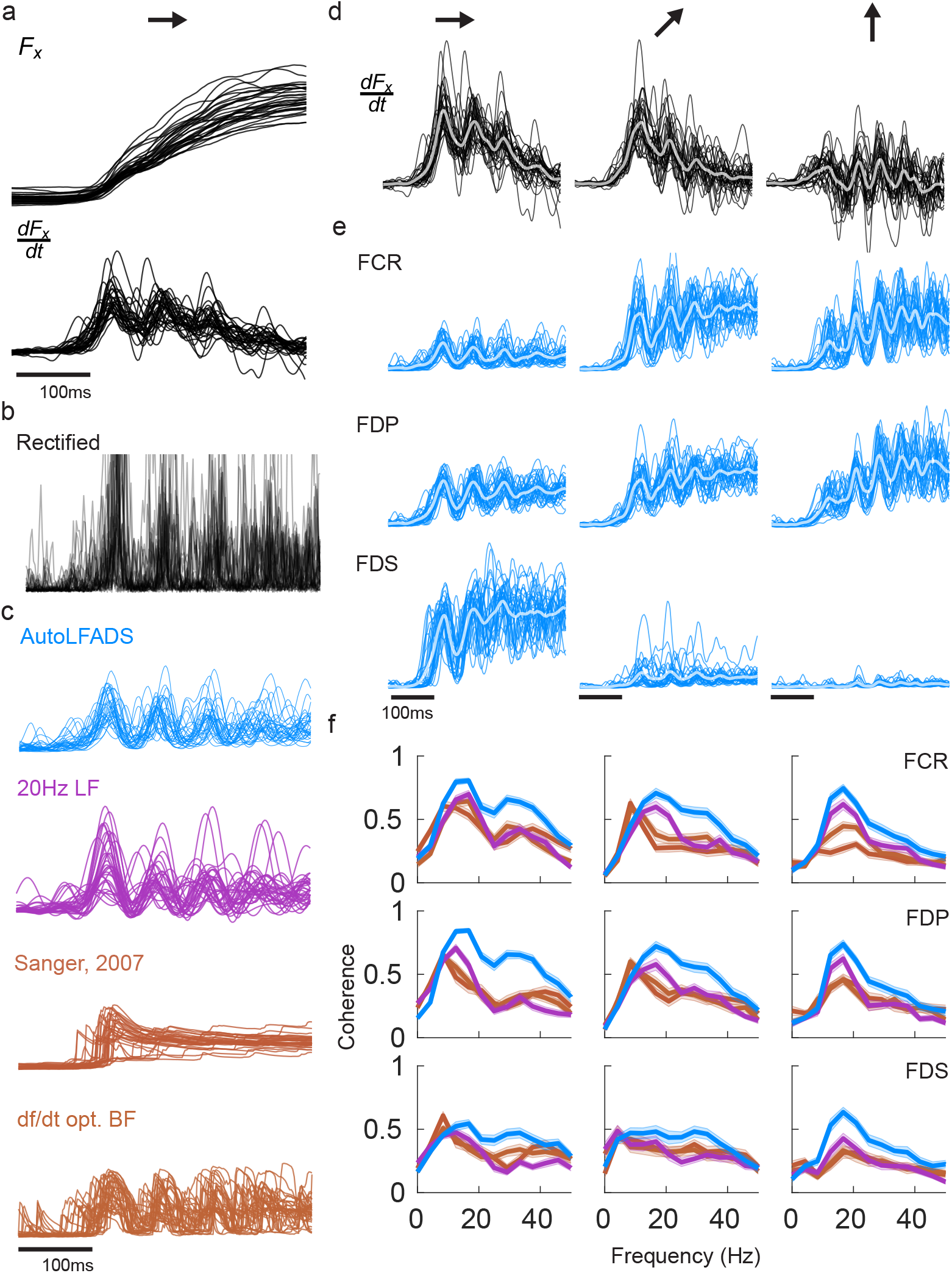
AutoLFADS enhances coherence to dF/dt. (a) Single trial oscillations were captured in the x component of recorded force (and more clearly in in dF_x_/dt) for the rightward target direction. (b) Corresponding oscillations were faintly visible in the rectified EMG. (c) Single trial muscle activation estimates from different approaches (d) Single trial oscillations of dF_x_/dt for three of the eight conditions, specifically to targets in the upper/right quadrant. Thick traces: trial average. (e) Single trial oscillations were also visible in AutoLFADS output across the wrist and digit flexor muscles (FCR: flexor carpi radialis, FDP: flexor digitorum profundus, FDS: flexor digitorum superficialis) (f) Magnitude-squared coherence computed between single trial EMG for a given muscle and dF_x_/dt) and averaged across trials. Each line represents the mean +/− SEM coherence to dF_x_/dt) for a given muscle activation estimate. Coherence was compared between AutoLFADS, lf_20_ EMG, Bayesian filtering with hyperparameters optimized to maximize prediction of dF/dt, and Bayesian filtering using the same hyperparameters as Sanger, 2007.

To quantify this correspondence, we computed the coherence between the muscle activation estimates and dF_x_/dt separately for each condition on a single trial basis. If the high-frequency features conserved by AutoLFADS accurately reflect underlying muscle activation, then they should have a closer correspondence with behavior (i.e., dF_x_/dt) than those features that remain after smoothing or Bayesian filtering. Coherence in the range of 10-50 Hz was significantly higher for AutoLFADS than for the other tested approaches, while all three muscle activation estimates had similar coherence to dF_x_/dt below 10 Hz (**Fig. 6f**). These results further demonstrate the power of the AutoLFADS approach to capture high-frequency features in the muscle activation estimates with higher fidelity than previous approaches - without requiring any choice of cutoff frequency or hyperparameters *a priori*.

### AutoLFADS preserves information about descending cortical commands

Simultaneous recordings of motor cortical and muscle activity gave us the ability to test how accurately AutoLFADS could capture the influence of descending cortical signals on muscle activation. We trained linear decoders to map the different muscle activation estimates onto each of the 96 channels of smoothed motor cortical activity. Note that while the causal flow of activity is from M1 to muscles, we predicted in the opposite direction (muscles to M1) so that we had a consistent target (M1 activity) to assess the informativeness of the different muscle activation estimates. For each channel of M1 activity, we smoothed the observed threshold crossings using a Gaussian kernel with standard deviation of 35ms. We performed predictions in a window spanning 150 ms prior to 150 ms after force onset. For each channel, we found the optimal decoding lag between neural and muscle activity by sweeping in a range from 0 to 100ms (i.e., we allowed predicted neural activity to precede the muscle activity predictors by up to 100 ms). AutoLFADS significantly improved the accuracy of M1 activity prediction when compared to 20Hz low pass filtering (**Fig. 7**; mean percentage improvement: 12.4%, p=2.32e-16, one-sided Wilcoxon paired signed rank test). Similarly, AutoLFADS significantly improved M1 prediction accuracy when compared to Bayesian filtering using previously published hyperparameters (mean percentage improvement: 22.2%, p=4.58e-16, one-sided Wilcoxon paired signed rank test) or using hyperparameters optimized to predict behavior (mean percentage improvement: 16.9%, p=7.37e-16, one-sided Wilcoxon paired signed rank test). This demonstrates that AutoLFADS not only captures behaviorally-relevant features, but also preserves signals that reflect activity in upstream brain areas that are involved in generating muscle activation commands.

**Fig. 7.**
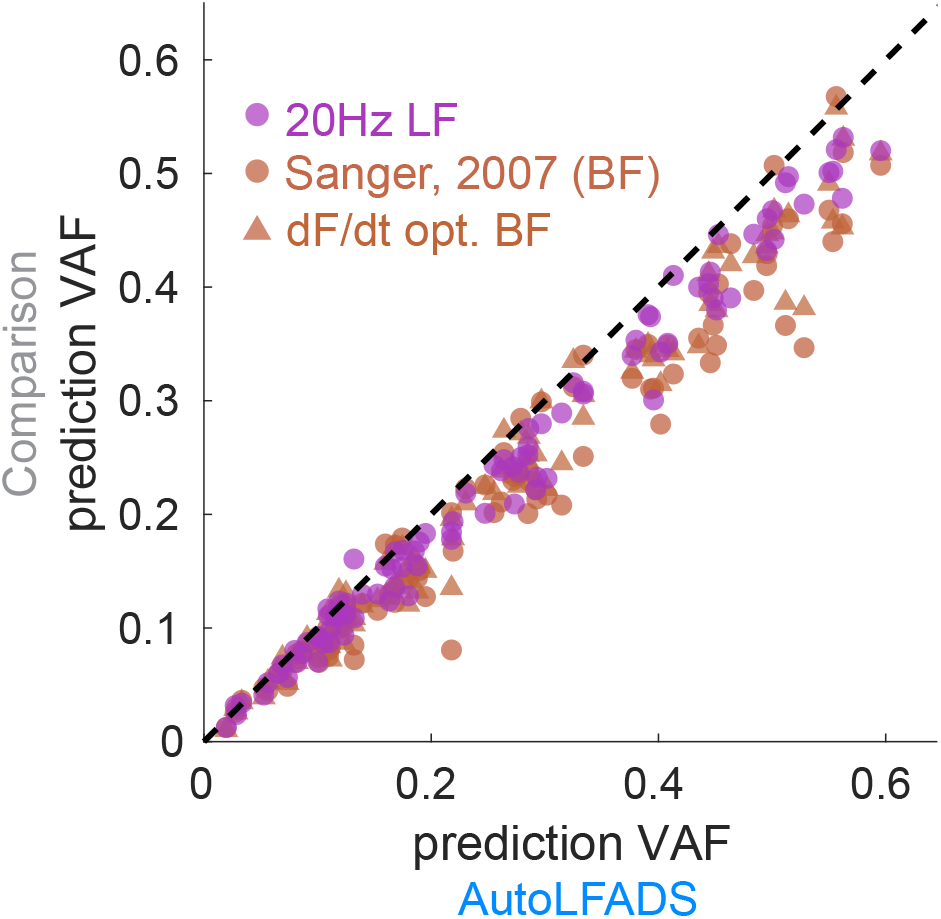
AutoLFADS is correlated with motor cortical (M1) activity. Scatter plot comparing accuracy of the prediction of motor cortical activity from EMG filtered using different approaches and AutoLFADS. Each dot represents prediction accuracy in terms of VAF for a single M1 channel.

## Discussion

We demonstrate that AutoLFADS provides a powerful tool to estimate muscle activation from EMG activity, complementing the analysis of cortical activity to which it has previously been applied. In our tests, AutoLFADS outperformed standard filtering approaches in estimating muscle activation. In particular, the AutoLFADS-inferred muscle activation commands were substantially more informative about both behavior and brain activity, all in an unsupervised manner that did not require manual tuning of hyperparameters. Further, we observed robust performance across data from two different species (rat and monkey) performing different behaviors that elicited muscle activations at varying timescales and with very different dynamics. The lack of prior assumptions about behaviorally-relevant timescales is critical, as there is growing evidence that precise timing plays an important role in motor control, and that high-frequency features may be important for understanding the structure of motor commands (Sober et al., 2018; Srivastava et al., 2017). We demonstrated that AutoLFADS is capable of extracting high-frequency features underlying muscle commands - such as muscle activity related to foot off during locomotion, or the oscillations uncovered during isometric force contraction - that are otherwise difficult to analyze using conventional filtering. By improving our ability to estimate short-timescale features in muscle activation, AutoLFADS may serve to increase the precision with which we analyze EMG signals when studying motor control.

### Relation to previous work

Substantial effort has been invested over the past 50 years to develop methods to estimate the latent command signal to muscles - termed the *neural drive* - through EMG recordings (Farina et al., 2014, 2004). This is a challenging problem as typical recordings reflect the summed activity of many muscle fibers in the vicinity of an electrode. This group of muscle fibers reflects the activity of potentially many different motor units, which can lead to cancellation of signal when summed together in recorded EMG (Negro et al., 2015). As muscle contraction increases, the linear relationship between the number of active motor units and the amplitude of the recorded EMG becomes distorted, making it non-trivial to design methods that can extract neural drive (Dideriksen and Farina, 2019).

Higher spatial resolution recordings may allow neural drive to be inferred with greater precision. For example, high-density EMG arrays have enabled access to the action potentials of as many as 30 to 50 individual motor units from the surface recordings of a single muscle (De Luca et al., 2006; Del Vecchio et al., 2020; Farina et al., 2016). By isolating the activity of individual motor units, one can better extract the common input signal within the fibers of a given muscle, a quantity that is more closely linked to force output than is the typical EMG (Boonstra et al., 2016; De Luca and Erim, 1994; Farina and Negro, 2015). Here, we showed that AutoLFADS similarly enhanced correspondence to force output and behavioral parameters better than standard techniques of EMG analysis; whether this improvement approaches that from using multiple motor unit recordings is not clear and is the goal of future work.

AutoLFADS leverages recent advances at the intersection of deep learning and neuroscience to create a powerful model that describes both the spatial and temporal regularities that underlie multi-muscle coordination. Previous work has attempted to estimate neural drive to muscles by exploiting either temporal regularities in EMGs using Bayesian filtering techniques applied to single muscles (Hofmann et al., 2016; Sanger, 2007) or spatial regularities in the coordination of activity across muscles (Alessandro et al., 2012; d’Avella et al., 2003; Hart and Giszter, 2004; Kutch and Valero-Cuevas, 2012; Ting and Macpherson, 2005; Torres-Oviedo and Ting, 2007; Tresch et al., 2006, 1999). Conceptually, AutoLFADS approaches the problem in a different way by using an RNN to simultaneously exploit spatial and temporal regularities across muscles. AutoLFADS builds on advances in training RNNs (e.g., using sequential autoencoders) that have enabled unsupervised characterization of complex dynamical systems, and large-scale optimization approaches that allow neural networks to apply out-of-the-box to a wide variety of data. Capitalizing on these advances allows a great deal of flexibility in estimating the regularities underlying muscle coordination. For example, our approach does not require an explicit estimate of dimensionality, does not restrict the modelled muscle activity to the span of a fixed time-varying basis set (Alessandro et al., 2012; d’Avella et al., 2003), does not assume linear interactions in the generative model underlying the observed muscle activations, and has an appropriate (and easily adapted) noise model to relate inferred activations to observed data. This flexibility enabled us to robustly estimate neural drive to muscles across a variety of conditions: different phases of the same behavior (stance vs. swing), different animals (rat vs. monkey), behaviors with distinct dynamics (locomotion vs. isometric contractions), and different time scales (100’s of ms vs. several seconds). AutoLFADS can therefore be used by investigators studying motor control in a wide range of experimental conditions, behaviors, and organisms.

### Limitations

EMG is susceptible to artifacts (e.g., powerline noise, movement artifact, cross-talk, etc.), therefore any method that aims to extract muscle activation from EMG necessitates intelligent choice of preprocessing to mitigate the potential deleterious effects of these noise sources (see Methods). As a powerful deep learning framework that is capable of modeling nonlinear signals, AutoLFADS has a specific sensitivity to even a small number of high-amplitude artifacts that can skew model training, since producing estimates that capture these artifacts can actually lower the reconstruction cost that the model is trying to minimize. Our implementation of data augmentation strategies during training helped mitigate the sensitivity of AutoLFADS to spurious high-magnitude artifacts. Our results showing the muscle activation estimates from AutoLFADS were more informative about behavior and brain activity than standard approaches, suggests that these pervasive noise sources in multi-muscle EMG recordings were not detrimental to the model’s performance on the datasets we tested. However, it is unclear whether this would be more of an issue in recordings with substantial noise. Future studies may be able to explore alternative methods, such as regularization strategies or weighting of reconstruction costs during training, to mitigate these effects of recording artifacts in addition to the data augmentation we developed in this study.

### Potential Future Applications

The applications of AutoLFADS may extend beyond scientific discovery to improved clinical devices that require accurate measurement of muscle activation. Current state of the art myoelectric prostheses use supervised classification to translate EMGs into control signals (Ameri et al., 2018; Hargrove et al., 2017; Vu et al., 2020). AutoLFADS could complement current methods through *unsupervised pre-training* to estimate muscle activation from EMG signals before they are input to classification algorithms. The idea of unsupervised pre-training has gained popularity in the field of natural language processing (Qiu et al., 2020). In these applications, neural networks that are pre-trained on large repositories of unlabeled text data uncover more informative representations than do standard feature selection approaches, which improves accuracy and generalization in subsequent tasks. Similar advantages have been demonstrated for neural population activity by unsupervised pre-training using LFADS, where the model’s inferred firing rates enable decoding of behavioral correlates with substantially higher accuracy and better generalization to held-out data than standard methods of neural data pre-processing (Keshtkaran et al., 2021; Pandarinath et al., 2018).

Designing AutoLFADS models that either don’t require - or require minimal - network re-training to work across subjects could help drive translation to clinical applications. Another area of deep learning research - termed *transfer learning* - has produced methods that transfer knowledge from neural networks trained on one dataset (termed the *source domain*) to inform application to different data (termed the *target domain*) (Zhuang et al., 2020). Developing subject-independent AutoLFADS models using transfer learning approaches would reduce the need for subject-specific data collection that may be difficult to perform outside of controlled, clinical environments. Subject-independent models of muscle coordination may also benefit brain-controlled functional electrical stimulation (FES) systems that rely on mapping motor cortical activity onto stimulation commands to restore muscle function (Ajiboye et al., 2017; Ethier et al., 2012). Components of subject-independent AutoLFADS models trained on muscle activity from healthy patients, could be adapted for use in decoder models that aim to map simultaneous recordings of motor cortical activity onto predictions of muscle activation for patients with paralysis. These approaches may provide a neural network solution to decode muscle activation commands from brain activity more accurately, serving to improve the estimates of stimulation commands for FES implementations. Combining AutoLFADS with transfer learning methods could also enable the development of new myoelectric prostheses that infer the function of missing muscles by leveraging information about the coordinated activation of remaining muscles.

There are several aspects of the AutoLFADS approach that might provide additional insights into the neural control of movement. First, AutoLFADS also provides an estimate of the time-varying input to the underlying dynamical system (i.e., **u**[t]). The input represents an unpredictable change to the underlying muscle activation state, which likely reflects commands from several upstream brain areas that converge with sensory feedback at the level of the spinal cord. Analyzing **u**[t] in cases where we also have simultaneous recordings of relevant upstream brain areas may allow us to distinguish the contributions of these brain and peripheral sources to muscle coordination, including the contributions of different reflex pathways (Pruszynski et al., 2011). Second, AutoLFADS may also be useful for studying complex, naturalistic behavior. In this study we applied AutoLFADS to activity from two relatively simple, yet substantially different motor tasks, each with stereotyped structure. However, because the method is applied to windows of activity without regard to behavior, it does not require repeated trials of the same movement or alignment to behavioral or task events. Even with this unsupervised approach, AutoLFADS preserved information about behavior (kinematics and forces) with higher fidelity than standard filtering that used supervision to preserve behaviorally-relevant features. As supervision becomes increasingly challenging during behaviors that are less constrained or difficult to carefully monitor (e.g., free cage movements), unsupervised approaches like AutoLFADS may be necessary to extract precise estimates of muscle activity. AutoLFADS may therefore have the potential to overcome traditional barriers to studying neural control of movement by allowing investigators to study muscle coordination during unconstrained, ethologically-relevant behaviors rather than the constrained, artificial behaviors often used in traditional studies.

## Acknowledgements

This work was supported by the Emory Neuromodulation and Technology Innovation Center (ENTICe), NSF NCS 1835364, DARPA PA-18-02-04-INI-FP-021, NIH Eunice Kennedy Shriver NICHD K12HD073945, the Alfred P. Sloan Foundation, the Burroughs Wellcome Fund, and the Simons Foundation as part of the Simons-Emory International Consortium on Motor Control (CP), NSF NCS 185345, NIH NINDS NS086973, NIH NINDS NS053603 and NS074044 (LEM), NIH NINDS NS086973 (MCT), UKRI EPSRC EP/T020970/1, Community of Madrid Talent Attraction Fellowship 2017-T2/TIC-5263 (JAG), German Academic Scholarship Foundation Fellowship 289999 and Elite Network of Bavaria Travel Allowance (JFB).

## Code availability

Code will be made available upon publication.

## Data availability

Data will be made available upon publication.

## Author Contributions

L.N.W. and C.P. designed the study, performed analyses, and wrote the manuscript with input from all other authors. L.N.W., M.R.K., and C.P. developed the algorithmic approach. J.F.B. contributed to analysis of rat data. D.H. contributed to application of Bayesian filtering. C.A. and M.C.T. recorded the data from the rat. J.A.G. and L.E.M. recorded data from the monkey. All authors contributed to manuscript revision.

## Methods

### Rat locomotion data

#### EMG recordings

Chronic bipolar EMG electrodes were implanted into the belly of 12 hindlimb muscles during sterile surgery as described in (Alessandro et al., 2018). Recording started at least two weeks after surgery to allow the animal to recover. Differential EMG signals were amplified (1000X), band-pass filtered (30–1000 Hz) and notch filtered (60 Hz), and then digitized (5000 Hz).

#### Kinematics

A 3d motion tracking system (Vicon) was used to record movement kinematics (Alessandro et al., 2020, 2018). Briefly, retroreflective markers were placed on bony landmarks (pelvis, knee, heel and toe) and tracked during locomotion at 200 Hz. The obtained position traces were low-pass filtered (Butterworth 4th order, cut-off 40 Hz). The 3-D knee position was estimated by triangulation using the lengths of the tibia and the femur in order to minimize errors due to differential movements of the skin relative to the bones (Bauman and Chang, 2010; Filipe et al., 2006). The 3d positions of the markers were projected into the sagittal plane of the treadmill, obtaining 2d kinematics data. Joint angles were computed from the positions of the top of the pelvis (rostral marker), the hip (center of the pelvis), the knee, the heel and the toe as follows. Hip: angle between the pelvis (line between pelvis-top and hip) and the femur (line between hip and knee). Knee: angle between the femur and the tibia (line between the knee and the heel). Ankle: angle between the tibia the foot (line between the heel and the toe). All joint angles are equal to zero for maximal flexion and increase with extension of the joints.

Joint angular velocities and accelerations were computed from joint angular position using a first-order and second-order Savitzky-Golay (SG) differentiation filter (MATLAB’s *sgolay*), respectively. SG FIR filter was designed with a 5^th^ order polynomial and frame length of 27.

### Monkey isometric data

#### M1/EMG Recordings

We recorded data from one 10 kg male Macaca mulatta monkey (age 8 years when the experiments started) while he performed an isometric wrist task. All surgical and behavioral procedures were approved by the Animal Care and Use Committee at Northwestern University. The monkey was implanted with a 96-channel microelectrode silicon array (Utah electrode arrays, Blackrock Microsystems, Salt Lake City, UT) in the hand area of M1, which we identified intraoperatively through microstimulation of the cortical surface. The monkey was also implanted with intramuscular EMG electrodes in a variety of wrist and hand muscles. We report data from the following muscles: flexor carpi radialis, flexor carpi ulnaris, flexor digitorum profundus (2 channels), flexor digitorum superficialis (2 channels), extensor digitorum communis, extensor carpi ulnaris, extensor carpi radialis, palmaris longus, pronator teres, abductor polis brevis, flexor polis brevis, and brachioradialis. Surgeries were performed under isoflurane gas anesthesia (1–2%) except during cortical stimulation, for which the monkeys were transitioned to reduced isoflurane (0.25%) in combination with remifentanil (0.4 μg kg^−1^ min^−1^ continuous infusion). Monkeys were administered antibiotics, anti-inflammatories, and buprenorphine for several days after surgery.

#### Experimental Task

Monkeys were trained at a 2D isometric wrist task for which the relationship between muscle activity and force is relatively simple and well characterized. Each monkey’s left arm was positioned in a splint so as to immobilize the forearm in an orientation midway between supination and pronation (with the thumb upwards). A small box was placed around the monkey’s open left hand, incorporating a 6 DOF load cell (20E12, 100 N, JR3 Inc., CA) aligned with the wrist joint. The box was padded to comfortably constrain the monkey’s hand and minimize its movement within the box. The monkeys controlled the position of a cursor displayed on a monitor by the force they exerted on the box. Flexion/extension force moved the cursor right and left, respectively, while forces along the radial/ulnar deviation axis moved the cursor up and down. Prior to placing the monkey’s hand in the box, the force was nulled in order to place the cursor in the center target. Being supported at the wrist, the weight of the monkey’s hand alone did not significantly move the cursor when the monkey was at rest. Targets were displayed either at the center of the screen (zero force), or equally spaced along a ring around the center target. All targets were squares with 4 cm sides.

### Data preprocessing

Raw EMG data was first passed through a powerline interference removal filtering algorithm (Keshtkaran et al., 2014) that we specified to remove power at 60Hz and 2 higher frequency harmonics (120Hz, 180Hz). We added an additional notch filtering step centered at 60Hz and 94Hz (isometric data) and 60Hz, 120Hz, 240Hz, 300Hz, and 420Hz (locomotion data) to remove noise peaks identified from observing the power spectral density estimates after the previous filtering step. We then filtered the EMG data with a 4th-order Butterworth high pass filter with cutoff frequency at 65 Hz. We finally rectified the EMG activity by taking the absolute value of the filtered EMG data. Rectified EMG was then down-sampled from the original sampling rate (rat: 5KHz, monkey: 2KHz) to 500Hz using MATLAB’s *resample* function which applies a poly-phase anti-aliasing filter prior to down-sampling the data.

To prepare the rectified EMG data for analysis, we next removed infrequent electrical recording artifacts that caused large spikes in activity. Removing artifactual signals is important for any analysis, but these had a particular effect on AutoLFADS modeling: during the modeling process (see *Training*), large anomalous events incurred a large reconstruction cost that disrupted the optimization process and prevented the model from capturing underlying signal. To remove artifacts, we clipped each channel’s activity such that any activation above a threshold percentile value was equalized to the threshold percentile value. For the isometric data, we applied the same 99.99^th^ percentile threshold across all channels. For the locomotion data, a 99.9^th^ percentile threshold was used for all channels except for one muscle (semimembranosus) which required a 99^th^ percentile threshold to clip multiple high-magnitude artifacts only observed in the recordings for this channel. After clipping, we normalized each channel by the 95th percentile value to ensure that all channels had comparable variances. We also scaled each channel’s activity by a factor of 1.5. Normalization was motivated by the assumption that the voltage amplitude of a given channel was not meaningful and likely reflected recording conditions for that particular electrode.

For the monkey isometric EMG data, some artifacts remained even after clipping. To more thoroughly reject artifacts, we extracted large windows of data that were aligned to force onset for successful trials (1.5s prior, 2.5s after force onset) and visually screened all muscles for anomalous events. Specifically, trials in which muscle activity contained high magnitude events that were not seen consistently across trials were excluded for subsequent analyses. After the rejection process, 227 of 296 successful trials remained.

We used AutoLFADS to model the EMG data without regard to behavioral structure (i.e., without regard to task or trial information). For the rodent locomotion data, the continuously recorded data (2 min) was binned at 2ms and divided into windows of length 200ms, with 40ms of overlap between the segments. The start indices of each trial are stored so that after training, model output can be reassembled. After performing inference on each segment separately, the segments were merged at overlapping regions using a linear combination of the estimates from the end of the earlier segment with those from the beginning of the later segment. The merging technique weighted the ends of the segments as *w* = 1 – *x* and the beginning of segments as 1 – *w*, with x linearly increasing from 0 to 1 across the overlapping points. After weights were applied, the overlapping segments were summed to reconstruct the modeled data. Non-overlapping portions of the windows were concatenated to the array normally. Due to the additional artifact rejection process for the monkey isometric data (mentioned above), we were unable to model the continuously recorded data (i.e., we could only model segments corresponding to trials that passed the artifact rejection process). These trial segments were processed using the same chopping process we administered to the rodent locomotion data.

### AutoLFADS architecture

The AutoLFADS model operates on segments of multi-channel rectified EMG data (**m**[t]). The output of the model 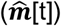 is an estimate of muscle activation from **m**[t] in which the observed activity on each channel is modeled as a sample from an underlying time-varying gamma distribution. For channel *j*, 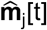 is the time-varying mean of the gamma distribution inferred by the model (i.e., AutoLFADS output). The two parameters for each gamma distribution, denoted *α* (concentration) and *β* (rate), are taken as separate linear transformations of the underlying dynamic state **s**[t] followed by an exponential nonlinearity. To generate **s**[t] for a given window of data **m**[t], a series of recurrent neural networks (RNNs), implemented using Gated Recurrent Units (GRU) cells, were connected together to explicitly model a dynamical system. Two RNNs (the Initial Condition & Controller Input Encoders) first operate on **m**[t] to generate an estimate of the initial state **s**[0] and to encode information for a separate RNN whose role is inferring a set of time-varying inputs into the Generator, **u**[t] (Controller). A learned dynamical system (Generator) models *f* to evolve **s**[t] at each timepoint. From **s**[0], the dynamical system is run forward in time with the addition of the **u**[t] at each timestep to evolve **s**[t].

The Generator, Controller, and Controller Input Encoder RNNs each had 64 units. The Initial Condition Encoder RNN had a larger dimensionality of 128 units to increase the capacity of the network to handle the complex mapping from the unfiltered rectified EMG onto the parameters of the initial condition distribution. The Generator was provided with a 30-dimensional initial condition (i.e., **s**[0]) and a three-dimensional input (i.e., **u**[t]) from the Controller. A linear readout matrix restricted the output of the Generator to describe the patterns of observed muscle activity using only 10 dimensions. Variance of the initial condition prior distribution was 0.1. The minimum variance of the initial condition posterior distribution was 1e-4.

### AutoLFADS training

Training AutoLFADS models on rectified EMG data largely followed previous applications of LFADS (Keshtkaran et al., 2021; Keshtkaran and Pandarinath, 2019; Pandarinath et al., 2018; Sussillo et al., 2016), with key distinctions being the gamma emissions process chosen for EMG data, as described above, and a novel data augmentation strategy implemented during training (see below). Beyond this choice, model optimization followed previous descriptions; briefly, the model’s objective function is defined as the log likelihood of the observed EMG data (**m**[t]) given the time-varying gamma distributions output by the model for each channel 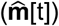, marginalized over all latent variables. This is optimized in a variational autoencoder by maximizing a lower bound on the marginal data log-likelihood (Kingma and Welling, 2014). Network weights were randomly initialized. During training, network parameters are optimized using stochastic gradient descent and backpropagation through time. We scaled the loss prior to computing the gradients (1e4) to prevent numerical precision issues. We used the Adam optimizer (epsilon: 1e-8, beta1: 0.9, beta2: 0.99) to control weight updates, and implemented gradient clipping to prevent potential exploding gradient issues (max. norm=200). To prevent any potentially problematic large values in RNN state variables and achieve more stable training, we also limited the range of the values in GRU hidden state by clipping values greater than 5 and lower than −5.

Finding a high-performing model required finding a suitable set of model hyperparameters (HPs). To do so, we used an automated model training and hyperparameter framework we developed (Keshtkaran and Pandarinath, 2019), which is an implementation of the population based training (PBT) approach (Jaderberg et al., 2017). Briefly, PBT is a large-scale optimization method that trains many models in parallel with different HPs. PBT trains models for a generation - i.e., a user-settable number of training steps - then performs a selection process to choose higher performing models and eliminate poor performers. Selected high performing models are also “mutated” (i.e., their HPs are perturbed) to expand the HP search space. After many generations, the PBT process converges upon a high performing model with optimized HPs.

For both datasets, we used PBT to simultaneously train 40 LFADS models for 30 epochs per generation (where an “epoch” is a single training pass through the entire dataset). We distributed training across 20 GPUs (2 models were trained on each GPU simultaneously). The magnitude of the KL and L2 penalties was linearly ramped for the first 50 epochs of training during the first generation of PBT. Each training epoch consisted of ~10 training steps. Each training step was computed on batches from the training data (monkey: 400 samples, rat: 400 samples). PBT stopped when there was no improvement in performance after 10 generations. Runs typically took ~30-40 generations to complete. After completion, the model with the lowest validation cost (i.e., best performance) was returned and used as the AutoLFADS model for each dataset. AutoLFADS output was inferred 200 times for each model using different samples from initial condition and controller output posterior distributions. These estimates were then averaged, resulting in the final model output used for analysis.

PBT was allowed to optimize the model’s learning rate and five regularization HPs: L2 penalties on the Generator and Controller RNNs, scaling factors for the KL penalties applied to the initial conditions and time-varying inputs to the Generator RNN, and two dropout probabilities (see *PBT Search Space* below for specific ranges).

A key distinction of the adapted AutoLFADS approach for EMG data with respect to previous application of LFADS, was the development and use of data augmentation strategies during training. In our initial modeling, we found that models could overfit to high-magnitude events that occurred simultaneously across multiple channels. To prevent overfitting to these sharp features in the data we implemented a novel data augmentation strategy (“temporal shift”). During model training, we randomly shifted the indexing of each channel for the windows of data input to the model. The model input and the data used to compute reconstruction cost were shifted differently to prevent the network from overfitting by drawing a direct correspondence between specific values of the input and output. During temporal shifting, the indexing of each channel was allowed to shift by a maximum of 6 bins. Shift values were randomly drawn from a normal distribution (stddev.= 3 bins) that was truncated such that any randomly drawn shift value greater than 6 was discarded and re-drawn. Finally, because rectified EMG follows a skewed distribution (i.e., values are strictly positive), data were log transformed before being input to the model, resulting in data distributions that were more symmetric.

#### PBT Search Space

During PBT, we specified search ranges (min/max) for regularization HPs. Weights were used to control maximum and minimum perturbation magnitudes for different HPs (e.g. a weight of 0.3 results in perturbation factors between 0.7 and 1.3). Keep ratio, dropout probability, and learning rate were limited to their specified ranges, while the KL and L2 penalties could be perturbed outside of the initial ranges.

| hyperparameter | min | max | perturbation weight | initial value |
| --- | --- | --- | --- | --- |
| learning rate | 1e-4 | 5e-3 | 0.3 | 2e-3 |
| keep ratio | 0.3 | 0.99 | 0.3 | 0.5 |
| L2 on generator | 1e-5 | 1e-1 | 0.8 | log-random |
| L2 on controller | 1e-5 | 1e-1 | 0.8 | log-random |
| KL weight on controller outputs | 1e-6 | 1e-4 | 0.8 | log-random |
| KL weight on initial conditions | 1e-6 | 1e-4 | 0.8 | log-random |
| dropout probability | 0.3 | 1 | 0.3 | random |

### Bayesian filtering

#### Model

Bayesian filtering is a nonlinear, adaptive filtering method that has previously been applied to surface EMG signals (Dyson et al., 2017; Hofmann et al., 2016; Sanger, 2007). A Bayesian filter consists of two components: (1) the temporal evolution model governing the time-varying statistics of the underlying latent variable and (2) the observation model that relates the latent variable to the observed rectified EMG. For this application, the temporal evolution model was implemented using a differential Chapman Kolmogorov equation with a diffusion term and a jump process (Hofmann et al., 2016). For the observation model, we tested both a Gaussian distribution and Laplace distribution, which are typical for application to surface EMG (Nazarpour et al., 2013). Empirically, we found that the Gaussian distribution produced slightly better decoding predictions (for both rat and monkey) than the Laplace distribution, so we used this model for all comparisons. Our implementation of the Bayesian filter had four hyperparameters: (1) sigma_max_, the upper interval bound for the latent variable, (2) number of bins used to define the histogram modeling the latent variable distribution, (3) alpha, which controls the diffusion process underlying the time-varying progression of the latent variable and (4) beta, which controls the probability for larger jumps of the latent variable. Sigma_max_ was empirically derived to be approximately an order of magnitude larger than the standard deviation measured from the aggregated continuous data. This value was set to 2 for both datasets after quantile normalizing the EMG data prior to modeling. The number of bins was set to 100 to ensure the changes in the magnitude of the latent variable over time could be modeled with high enough resolution. The other two hyperparameters, alpha and beta, had a significant effect on the quality of modeling, therefore we performed a large-scale grid search to optimize the parameters for each dataset (described below). Each EMG channel is estimated independently using the same set of hyperparameters.

#### Hyperparameter optimization

Two hyperparameters (alpha and beta) can be tuned to affect the output of the Bayesian filter. To ensure that the choice to use previously published hyperparameters (Sanger, 2007) was not resulting in sub-optimal estimates of muscle activation, we performed grid searches for each dataset to find an optimal set of hyperparameters to filter the EMG data. We tested 4 values of alpha (ranging from 1e-1 to 1e-4) and 11 values of beta (ranging from 1e0 to 1e-10), resulting in 44 hyperparameter combinations. To assess optimal hyperparameters for each dataset, we fit linear decoders from the Bayesian filtered EMG to predict joint angular acceleration (rat locomotion) or dF/dt (monkey isometric) following the same procedures used for decoding analyses (see *Predicting joint angular acceleration* or *Predicting dF/dt* for details). The hyperparameter combination that resulted in the highest average decoding performance across prediction variables was chosen to be optimal for comparison to AutoLFADS. Optimal hyperparameters were alpha=1e-2 and beta=1e-4 (rat locomotion) and alpha=1e-2 and beta=1e-2 (monkey isometric).

### Analyses

#### Metric

To quantify decoding performance, we used Variance Accounted For (VAF). 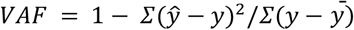. 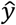 is the prediction, *y* is the true signal, and 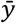 is the mean. This is often referred to as the coefficient of determination, or R^2^.

#### Predicting joint angular acceleration (rat locomotion)

We used optimal linear estimation (OLE) to perform single timepoint decoding to map the muscle activation estimates of 12 muscles (12 channels) onto the 3 joint angular acceleration kinematic parameters (hip, knee, and ankle). We performed a 10-fold cross validation (CV) to fit decoders (90% of the data) and evaluate performance (held-out 10% of data). Mean and standard error of the mean (SEM) for decoding performance was reported as the averaged cross-validated VAF across the 10 folds. During fitting of the decoders, we normalized the predictor (i.e., muscle activation estimate) and response (i.e., joint angular kin.) variables to have zero-mean and unit standard deviation. We also penalized the weights of the decoder using an L2 ridge regularization penalty of 1. We assessed significance using paired t-tests between the cross-validated (CV) VAF values from the AutoLFADS predictions and the predictions from the best performing low pass filter (10 CV folds). We performed a sweep to find the optimal lag ranging from 0ms to 200ms for each predictor with respect to each response variable. This allowed different muscle activation estimates to optimize their decoding performance by allowing for different lag relationships with respect to different joint angular kinematic parameters in the locomotion task. Optimal lag was determined for each muscle activation estimate and each response variable based on the highest mean CV VAF value.

#### Estimating frequency responses

We estimated frequency responses in order to understand how AutoLFADS acts differently than the typical approach of low pass filtering with a Butterworth filter, which applies the same temporally invariant frequency response to all channels. The intuition behind this analysis is to see whether AutoLFADS differently treats EMG signals at different characteristic timepoints and from different muscles. The magnitude of the frequency response of a linear dynamical system can be computed by dividing the power spectrum of its output by the power spectrum of its input. AutoLFADS is a highly non-linear system, but the linear frequency response still provides an interpretable measurement of the system’s behavior.

We chose two characteristic time points within the gait cycle of the rat locomotion dataset: (1) during the swing phase (at ¼ of the time between foot off and foot strike) and (2) during the stance phase (in the middle between foot off and foot strike). We selected windows of 175 samples (350 ms) around both of these time points. The number of samples was optimized to be large enough to have a fine resolution in frequency space but small enough to mainly include samples from one characteristic phase of the cycle (avg. stance: ~520 ms; avg. swing: ~155 ms). To reduce the effects of frequency resolution limits imposed by estimating PSDs on short sequences, we used a multi-taper method that matches the length of the windows used for the PSD estimates to the length of the isolated segment. This maximized the number of points we use to describe the frequency spectra of the isolated segments without upsampling. We z-scored the signals within each cycle and muscle to be able to compare frequency spectra across muscles and between different time points of the gait cycle. Within each window, we computed the Thomson’s multitaper power spectral density (PSD) estimate using a MATLAB (Mathworks Inc, MA) built-in function *pmtm*. We set the time-bandwidth product to NW=3 to retain precision in frequency space and the number of samples for the DFT to equal the window length and otherwise retained default parameters. With a window of 175 samples, the DFT has a frequency resolution of 2.85 Hz. Calculating the multi-taper estimate of the PSD with NW=3 lowers the resolution to 17.1 Hz.

We computed the PSD for the rectified EMG (*PSD_raw_*), the output of AutoLFADS (*PSD_AutoLFADS_*) and the output of a 20 Hz Gaussian low pass filter (*PSD*_*lf*_20__) within each window. We then computed the linear frequency response of AutoLFADS as *FR_AutoLFADS_* = *PSD_AutoLFADS_* / *PSD_raw_* and for the low pass filter as *FR*_*lf*_20__ = *PSD*_*lf*_20__ / *PSD_raw_*. Finally, we computed the mean and SEM across gait cycles. This analysis yielded frequency responses at two different time points in the gait cycle for every muscle.

#### Predicting dF/dt (monkey isometric)

To compute force onset, dF/dt was isolated in windows around target onset (250ms prior, 750ms after) for successful trials. Force onset was determined as the threshold crossing point for 15% of the max dF/dt in the isolated window. We used OLE to perform single timepoint decoding to map the EMG activity of 12 muscles (14 channels) onto the 2 components (i.e., X and Y) of dF/dt. We performed the same CV scheme used in the rat locomotion joint angular decoding analysis to evaluate decoding performance of the different EMG representations. During fitting of the decoders, we normalized the predictor variables (i.e., EMG) to have zero-mean and unit standard deviation. We penalized the weights of the decoder using an L2 ridge regularization penalty of 1.

#### Predicting M1 activity

We used OLE to perform single timepoint linear decoding to map the EMG activity of 12 muscles (14 channels) onto the 96 channels of Gaussian smoothed motor cortical threshold crossings (standard deviation of Gaussian kernel: 35ms). We performed the same CV scheme used in the rat locomotion joint angular decoding analysis to evaluate decoding performance for prediction of the inputs from motor cortical activity. During fitting of the decoders, we normalized the predictor variables (i.e., EMG) to have zero-mean and unit standard deviation. We penalized the weights of the decoder using an L2 ridge regularization penalty of 1. We performed a sweep for the optimal lag (ranging from 0 to 150ms) for each M1 channel. Optimal lag was determined in the same manner used for the EMG decoding analyses. To assess significance, we performed a Wilcoxon signed-rank test using MATLAB command *signrank* to perform a one-sided nonparametric paired difference test comparing the mean CV VAF values for predictions of all 96 channels.

#### Oscillation analyses

To improve alignment of oscillations observed in AutoLFADS output from the monkey isometric task, we used an algorithm that optimizes the lead or lag for each single trial in order to minimize the residual between the single trial activity and the trial-average for a given condition (Williams et al., 2020). We provided the algorithm with a window of AutoLFADS output around the time of force onset (80ms prior, 240ms after), and constrained the algorithm to limit the maximum shift of an individual trial to be approximately half the width of an oscillation (~35ms) and for the template to be non-negative. For all applications of the optimal shift algorithm, we applied a smoothness regularization scale of 1.0.

For coherence analyses, we isolated a window of EMG activity and dF/dt around the alignment from optimal shifting of the AutoLFADS output (300ms prior, 600ms after). For each condition, we used the MATLAB function *mscohere* to compute coherence between the single trial activity of a given muscle’s activity and the dF/dt component. We specified the power spectral density estimation parameters within *mscohere* to ensure a robust calculation on the single trial activity. We used Hanning windows with 120 timesteps for the FFT and window size, and 100 timesteps of overlap between windows.

